# Genetic variation in the Major Histocompatibility Complex and association with depression

**DOI:** 10.1101/469023

**Authors:** Kylie P Glanville, Jonathan R I Coleman, Ken B Hanscombe, Jack Euesden, Shing Wan Choi, Kirstin L Purves, Gerome Breen, Tracy M Air, Till F M Andlauer, Bernhard T Baune, Elisabeth B Binder, Douglas H R Blackwood, Dorret I Boomsma, Henriette N Buttenschøn, Lucía Colodro-Conde, Udo Dannlowski, Nese Direk, Erin C Dunn, Andreas J Forstner, Eco JC de Geus, Hans J Grabe, Steven P Hamilton, Ian Jones, Lisa A Jones, James A Knowles, Zoltán Kutalik, Douglas F Levinson, Glyn Lewis, Penelope A Lind, Susanne Lucae, Patrik K Magnusson, Peter McGuffin, Andrew M McIntosh, Yuri Milaneschi, Ole Mors, Sara Mostafavi, Bertram Müller-Myhsok, Nancy L Pedersen, Brenda WJH Penninx, James B Potash, Martin Preisig, Stephan Ripke, Jianxin Shi, Stanley I Shyn, Jordan W Smoller, Fabian Streit, Patrick F Sullivan, Henning Tiemeier, Rudolf Uher, Sandra Van der Auwera, Myrna M Weissman, Major Depressive Disorder Working Group of the Psychiatric Genomics Consortium, Paul F O’Reilly, Cathryn M Lewis

## Abstract

**Background:** The prevalence of depression is higher in individuals suffering from autoimmune diseases, but the mechanisms underlying the observed comorbidities are unknown. Epidemiological findings point to a bi-directional relationship - that depression increases the risk of developing an autoimmune disease, and vice-versa. Shared genetic etiology is a plausible explanation for the overlap between depression and autoimmune diseases. In this study we tested whether genetic variation in the Major Histocompatibility Complex (MHC), which is associated with risk for autoimmune diseases, is also associated with risk for depression.

**Method:** We fine-mapped the classical MHC (chr6: 29.6-33.1 Mb), imputing 216 Human Leukocyte Antigen (HLA) alleles and four Complement Component 4 (C4) haplotypes in studies from the Psychiatric Genomics Consortium (PGC) Major Depressive Disorder (MDD) working group and the UK Biobank (UKB). In the 26 PGC-MDD studies, cases met a lifetime diagnosis of MDD, determined by a structured diagnostic interview. In the UKB, cases and controls were identified from an online mental health questionnaire. The total sample size was 45,149 depression cases and 86,698 controls. We tested for association between depression status and imputed MHC variants in each study and performed an inverse-variance weighted meta-analysis across the PGC-MDD and UKB samples, applying both a conservative region-wide significance threshold (3.9-e6) and a candidate threshold (1.6e-4).

**Results:** No HLA alleles or C4 haplotypes were associated with depression at the conservative threshold in the PGC, UKB or meta-analysis. HLA-B*08:01 was associated with modest protection for depression at the candidate threshold in the meta-analysis. Under the conservative threshold, 70 SNPs were detected in the UKB and 143 SNPs were detected in the meta-analysis, mirroring previous findings from highly powered GWAS of depression.

**Discussion:** We found no evidence that HLA alleles, which play a major role in the genetic susceptibility to autoimmune diseases, or C4 haplotypes, which are strongly associated with schizophrenia, confer risk for depression. These results indicate that autoimmune diseases and depression do not share common risk loci of moderate or large effect in the MHC.

## Introduction

Depression is a debilitating psychiatric disorder with an estimated lifetime prevalence of 15%^1^, making it the leading cause of global disability^2^. The disorder is characterised by heterogeneous symptoms profiles^3^ and variable treatment outcomes^4^. Developing effective pharmaceutical treatments relies on uncovering the etiology of a disorder^5^, and the field of psychiatric genetics has made great progress toward this objective in the last decade by devoting substantial resources to the study of depression^6,7^. Despite this progress, the underlying biology of depression is still not fully understood. Comorbid psychiatric and physical traits may indicate shared biological pathways and provide a path to uncovering the etiology of idiopathic psychiatric disorders^8^. Here, we focus on comorbid autoimmune diseases and depression, consider the mechanisms that could drive the overlap, and test for evidence of shared genetic influences in the Major Histocompatibility Complex (MHC).

Epidemiological studies indicate that individuals with a history of autoimmune disease are at greater risk of developing mood disorders compared to individuals without a history of autoimmune disease^9–12^. For example, a Danish Registry study^9^ identified individuals who had hospital contact for autoimmune diseases or mood disorders over a thirty year period and showed that the risk of developing a mood disorder increased following onset of any autoimmune disease (Incident Rate Ratio (IRR) = 1.45; 95% CI = 1.39-1.52).

One interpretation is that the distress arising from autoimmune afflictions is causal to the onset of a mood disorder. However, other evidence indicates that the relationship is bi-directional^13,14^. For example, another Danish Registry study^13^ showed that individuals with a history of depression were at increased risk for developing any autoimmune disease, compared to those without a history of depression (IRR = 1.25, 95% CI = 1.19-1.31), and that this increase remained relatively stable across the first decade after diagnosis of depression.

There are a number of plausible explanations for the observed overlap between depression and autoimmunity. Shared environmental influences may increase risk for both disorders, e.g. stress is a risk factor for autoimmune disease^15^, and there is a phenotypic and genetic correlation between anxiety and depression^16^. Another view is that shared genetic influences act on autoimmune disease and depression through common immune pathways. Inflammation is a hallmark characteristic of autoimmune disease^17^ and elevated levels of pro-inflammatory cytokines have been observed in some individuals with depression^18^. Despite increasing interest in the role of inflammatory pathways in the pathogenesis of depression^19^, the role of genetic variation in the MHC, which plays a crucial role in human immunity^20^, has not been thoroughly interrogated in the context of depression.

The MHC is divided into three functionally distinct regions: class I and II regions contain highly polymorphic human leukocyte antigen (HLA) genes that are strongly associated with risk for autoimmune disease^17,21,22^, and the class III region contains complement component 4 genes (C4), which are strongly associated with risk for schizophrenia^23^. Three recent genome-wide association studies (GWAS) indicate that genetic variation within the MHC is involved in risk for depression. The largest GWAS of depression from the Psychiatric Genomics Consortium^24^ identified 44 independent loci associated with depression including a SNP in the classical class I region. A GWAS of a broadly defined depression phenotype in the UK Biobank^25^, revealed a peak of association in the MHC, with the strongest evidence for association also coming from the classical class I region. A recent meta-analysis^26^ combined summary statistics from the latter two studies^24,25^ with estimates from a GWAS of depression including 23andMe data^27^, and identified a SNP within the extended class I region with strong evidence for association with depression.

Highly polymorphic loci and long-range linkage-disequilibrium in the MHC complicate the interpretation of SNP associations^24^. However, with the advent of methods for imputing HLA alleles^28^ and C4 haplotypes^23^, there is an opportunity to dissect SNP signal in the region with fine-mapping techniques. We used this approach to test whether genetic variation associated with risk for autoimmune disease and schizophrenia is also associated with risk for depression. To our knowledge, this is the first study to leverage imputation to interrogate the involvement of HLA alleles and C4 haplotypes in depression. Our efforts should lead to a better understanding of the role of these loci in depression, and may provide insights into the mechanisms driving comorbid autoimmunity and depression.

## Methods

### Participants

Participant data came from a subset of the Major Depressive Disorder Working Group of the Psychiatric Genomics Consortium (PGC-MDD)^24^ and from the UK Biobank (UKB)^29^ to give a total of 131,847 individuals of European ancestry (55% female, 45,149 depression cases and 86,698 controls). We selected a subset of participants from the PGC-MDD study because individual-level genotype data, required for imputation of HLA alleles and C4 haplotypes, were not available for the entire PGC-MDD cohort. Individual-level genotype and phenotype data were available for 26 studies shared with the PGC, totaling 39,145 individuals (54% female, 15,805 cases, 23,340 controls). Across the PGC-MDD studies, structured diagnostic interviews were conducted to identify cases with a lifetime diagnosis of MDD according to the Diagnostic and Statistical Manual of Mental Health Disorders, 4^th^ Edition^30^, the International Classification of Diseases, 9^th^ Edition (ICD-9)^31^, the International Classification of Diseases, 10^th^ Edition (ICD-10)^32^ or the Composite International Diagnostic Interview Short Form (CIDI-SF)^33^. In most of the PGC-MDD studies, bipolar disorder, non-affective psychosis and substance use disorder were exclusion criteria in the cases, and controls were screened for absence of MDD and other psychiatric disorders. Ethical approvals were obtained by Principal Investigators of each study, with all participants giving full informed consent.

The UKB is a prospective cohort study that has collected genotype and phenotype data for over 500,000 individuals across the UK, aged between 40 and 69 at the time of recruitment^29^. 157,366 UKB participants completed an online Mental Health Questionnaire (MHQ) which assesses lifetime depressive disorder^34^. Using the recommended MHQ scoring protocol^34^, we identified 29,344 individuals with lifetime depressive disorder and 63,358 controls. Cases were excluded if they endorsed diagnosis of psychosis or bipolar disorder. Controls were excluded if they endorsed diagnosis of any psychiatric disorder in the MHQ, or self-reported depression or use of anti-depressant medication at baseline and follow-up interviews, or suffered from a mood disorder according to Hospital Episode Statistics, or met the criteria for a mood disorder according to Smith, et al.^35^. Further details of the PGC and UKB samples are in Supplementary Tables 1a and 1b. Access to data from the UKB was granted under application number 18177.

### Genotyping and quality control

Genotyping arrays used in the PGC and UKB samples are shown in Supplementary Table 1a. Quality control (QC) of genotype data in the 26 PGC studies was performed by the PGC Statistical Analysis Group using the ricopili pipeline^24^ such that variants and individuals below the following thresholds were retained: SNP missingness (before individual QC) < 0.05, individual missingness < 0.02, SNP missingness (after individual QC) < 0.02, deviation from heterozygosity |F_het_| < 0.20, HWE p-value > 10^-10^ (cases) and p-value > 10^-6^ (controls). After imputation with the 1000 Genomes reference panel^24^, SNPs with INFO > 0.8 and minor allele frequency (MAF) > 0.05 were retained for relatedness testing and principal component analysis. One individual from each pair with relatedness > 0.2 was removed and only individuals of European ancestry were retained for analysis.

Using genotype data that had undergone preliminary QC by the UKB^29^, we created an inclusion list of individuals of European ancestry using 4-means clustering on the first two principal components provided by the UKB. We created an exclusion list of individuals using relatedness kinship (KING) estimates provided by the UKB, and removed one individual from each pair up to 3rd-degree relationships (KING r^2^ > 0.044)^36^. In the remaining data we applied QC such that variants and individuals below the following thresholds were retained: SNP missingness (before individual QC) < 0.02, individual missingness < 0.02, SNP missingness (after individual QC) < 0.02, MAF > 0.01, HWE p-value > 10^-8^. The UKB^29^ imputed SNPs using the IMPUTE4 software^29^ with the Haplotype Reference Consortium (HRC) reference panel^37^ and the UK10K Consortium reference panel^38^ to produce dosage data in BGEN file format (version 1.2)^39^. We extracted imputed SNPs from the classical class I MHC (chr6: 29,640,000-33,120,000) and converted to PLINK 2 binary format (pgen, pvar, psam) for association analyses in PLINK 2.0^40^.

### HLA allele and C4 haplotype imputation

HLA alleles were imputed using genotype data from the PGC studies using the SNP2HLA software^28^ with the Type 1 Diabetes Genetics Consortium (T1DGC) reference panel^41^ to produce dosage data in Beagle format^42^. The T1DGC reference panel contains MHC haplotype information to enable imputation of HLA alleles at two-digit and four-digit resolution in eight HLA genes: *HLA-A, HLA-B* and *HLA-C* in the classical class I MHC and *HLA-DRB1, HLA-DQA1, HLA-DQB1, HLA-DPA1* and *HLA-DPB1* in the classical class II MHC.

HLA alleles were imputed in the UKB by the core analytical team using the HLA*IMP:02 software^29^ with multi-population reference panels^43^. Collectively, the reference panels contained MHC haplotype information to enable imputation of HLA alleles in 11 HLA genes: *HLA-A, HLA-B* and *HLA-C* in the classical class I MHC and *HLA-DRB5, HLA-DRB4, HLA-DRB3, HLA-DRB1, HLA-DQA1, HLA-DQB1, HLA-DPA1* and *HLA-DPB1* in the classical class II MHC. HLA alleles were encoded as biallelic in the PGC and UKB such that imputed dosages referred to the presence of 0, 1 or 2 copies of each HLA allele.

C4 haplotypes were imputed using genotype data in the PGC and UKB studies using the SNP2HLA software^28^ with the C4 reference panel developed by the McCarroll lab^41^ (http://mccarrolllab.com/wp-content/uploads/2014/12/MHC_haplotypes_CEU_HapMap3_ref_panel.bgl) to produce dosage data in Beagle format^42^. The reference panel consists of SNP and C4 haplotypes within the extended MHC (25 to 34 Mb on chromosome 6) for 110 individuals from the HapMap CEU population. The reference panel contains 17 C4 haplotypes, defined by copy number variation of *C4A* and *C4B* genes in short and long form (defined by a HERV insertion). Four C4 haplotypes with frequency > 0.01 were retained for analysis: AL-AL, AL-BL, AL-BS and BS (where A and B correspond to two isotypes of the C4 gene and L and S correspond to the long and short forms). Three of the common C4 haplotypes (AL-AL, AL-BL and AL-BS) segregate on two, three and five different SNP haplotypes, respectively. Association results for these C4 haplotypes were calculated by meta-analysing across SNP haplotypes corresponding to each C4 structure.

### Statistical analyses

In the PGC we tested each HLA allele and C4 haplotype for association with MDD case/control status using an additive logistic regression model applied to dosage data. We included 6 principal components (calculated by the PGC) to control for population structure. We extracted association results for SNPs in the classical class I MHC from PGC analyses in each study, applying a second layer of QC such that only variants with a minor allele frequency >0.01 and an info score ≥0.6 were retained. Post-QC variants were meta-analysed across the 26 PGC studies using an inverse-variance weighted approach.

In the UKB we tested each HLA allele, C4 haplotype and imputed SNP for association with depression case/control status using an additive linear regression model applied to dosage data. We regressed 6 principal components (calculated by the UKB), batch and centre on the depression phenotype using logistic regression in R 3.4.1^44^ and used the residuals as the outcome variable in subsequent linear regression. We then filtered on variants with a minor allele frequency above 0.01 and an info score ≥0.6 before meta-analysing across the PGC and UKB results. Analyses were performed using PLINK (version 1.9 and version 2.0)^40^.

To calculate the MHC region-wide significance threshold, we used the Genetic Type 1 Error Calculator^45^, an online resource that calculates the number of effective tests by estimating linkage disequilibrium (LD) between variants and applying a Bonferroni correction. We calculated a conservative region-wide significance threshold (3.9e-06), controlling for all imputed SNPs in the classical MHC, and a candidate significance threshold (1.6e-4), controlling only for HLA intragenic SNPs in the classical MHC.

The power calculation using the Genetic Power Calculator^39^ showed that with a sample size of 45,149 cases and 86,698 controls, for an HLA allele of frequency 0.05 (the median in our study), we had 80% power to detect an odds ratio above 1.09, at a region-wide significance level of alpha = 3.9e-06.

We compared the imputation accuracy and frequency of HLA alleles and C4 haplotypes in the PGC and UKB for variants present in both samples. The average imputation info score and frequency was calculated by weighting variant info scores and frequencies by the effective sample size in each PGC study calculated as 4 / (1/[number of cases] + 1/[number of controls])^40^.

Drawing on evidence from epidemiological studies^9,13^, we identified autoimmune diseases with evidence for a bi-directional relationship with depression. We conducted a literature review on PubMed using the search terms “HLA” and the relevant autoimmune disease to identify HLA risk alleles. We filtered on HLA alleles with evidence for independent association (p < 3.9e-06) in European populations. We evaluated evidence for involvement of these HLA alleles in depression, selecting those with MAF > 0.05 in our study and applying a Bonferroni correction to adjust for the number of HLA alleles investigated (0.05/15 = 0.003).

A variable for genetically predicted C4A brain expression was calculated for each individual in the PGC and UKB samples. We leveraged work from Sekar, et al., who fit a linear model between C4 structures and C4A brain expression measured in post-mortem brain tissue^23^. We used the betas from this model (provided in the supplementary materials by Sekar, et al.) to estimate C4A brain expression corresponding to each C4 haplotype present in the reference panel (Supplementary Table 2). Individual scores for C4A brain expression were calculated by multiplying the dosage for each C4 haplotype by the corresponding value for C4A brain expression. We tested genetically predicted C4A brain expression for association with depression case/control status in the PGC and UKB samples, controlling for six principal components. Genetically predicted C4A brain expression was also used as a covariate in conditional analyses on HLA alleles.

## Results

In total, 207 HLA alleles were imputed in at least two PGC studies, and 102 HLA alleles were imputed in the UKB, of which 93 were shared across data sets (Table 1). Variants imputed in either data set were included in the final meta-analysis (minimum effective sample size was 669 for HLA-B-3906 in the PGC). Four C4 haplotypes (AL-AL, AL-BL, AL-BS, BS) were imputed in all data sets.

**Table 1:** Variants imputed in at least two of the 26 PGC studies and the UKB.

| Gene | PGC | UKB | Variants in both PGC and UKB | Variants in either PGC or UKB |
| --- | --- | --- | --- | --- |
| *HLA-A* | 31 | 13 | 13 | 31 |
| *HLA-B* | 48 | 18 | 18 | 48 |
| *HLA-C* | 30 | 14 | 14 | 30 |
| *HLA-DPA* | 5 | 3 | 3 | 5 |
| *HLA-DPB* | 25 | 11 | 11 | 25 |
| *HLA-DQA* | 12 | 7 | 7 | 12 |
| *HLA-DQB* | 19 | 12 | 12 | 19 |
| *HLA-DRB* | 37 | 24 | 15 | 46 |
| **Total HLA alleles** | **207** | **102** | **93** | **216** |
| **C4 haplotypes** | **4** | **4** | **4** | **4** |
| **SNPs** | **49,611** | **47,799** | **40,561** | **56,779** |

There was strong consistency between the frequency and info scores of HLA alleles and C4 haplotypes imputed in both the PGC and UKB (correlation r=0.99 for frequency and r=0.86 for info score; Supplementary Figures 1 and 2). For the 93 HLA alleles imputed in both data sets, the info score was consistently higher in the UKB compared to the PGC studies (UKB mean = 0.98, PGC mean = 0.96), possibly due to the larger HLA reference panel, and/or greater efficiency of the imputation algorithm, used by the UKB.

Testing for association with depression in the PGC sample, no HLA allele, C4 haplotype or SNP surpassed region-wide significance (Figure 1a). In the UKB, no HLA allele or C4 haplotype surpassed region-wide significance (Figure 1b). The strongest evidence for association with an HLA allele was HLA-B*08:01 (p = 4e-4, OR = 0.98, 95% CI = 0.97-0.99). Among SNPs, 70 met region-wide significance (Supplementary Table 3). The variant with the lowest p-value (“UKB index variant”) was a SNP in the classical class I region; rs1264373 (p = 3.21e-7, OR = 0.97, 95% CI = 0.96-0.98). All variants surpassing region-wide significance in the UKB sample were in LD with the UKB index variant (0.66 < R^2^ < 1.00). The UKB index variant was also in LD with the most significant SNP within the MHC reported in the PGC MDD GWAS^24^ (rs115507122, R^2^ = 0.63).

**Figure 1:**
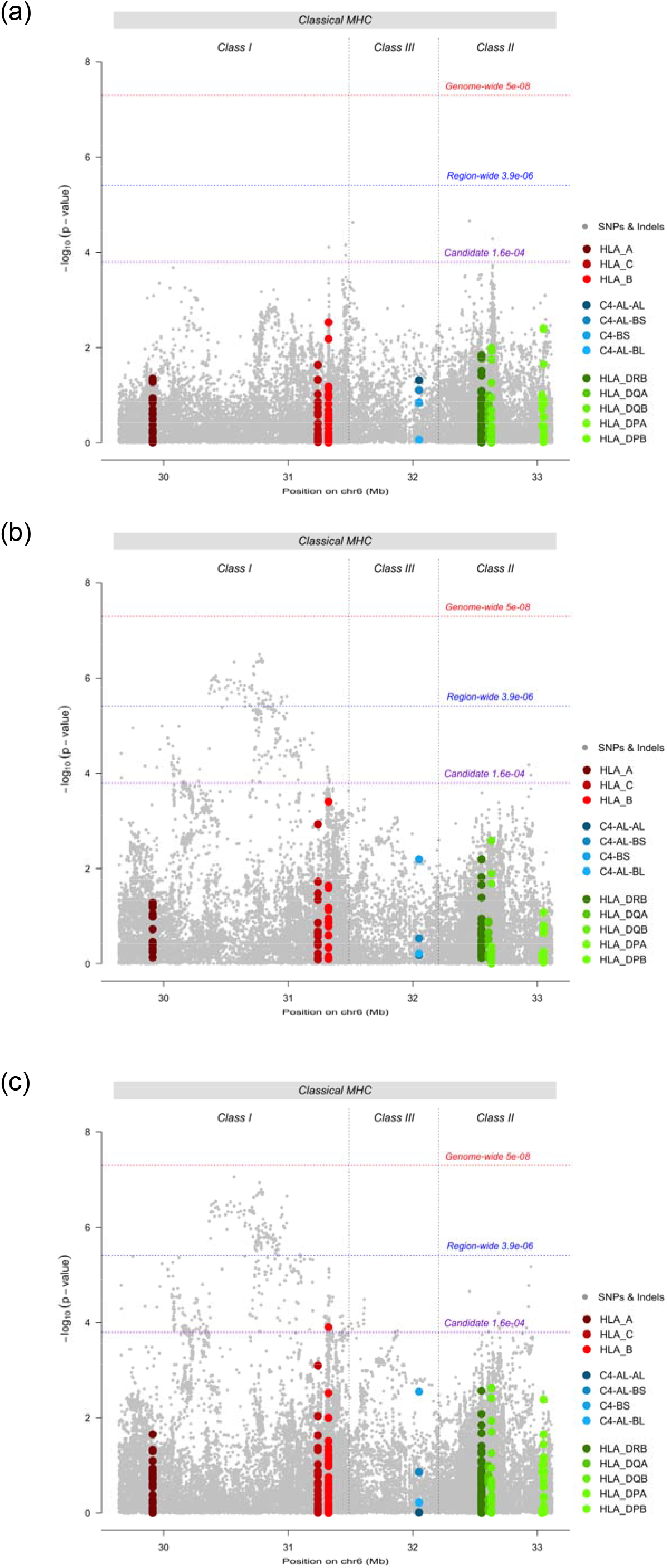
Region-wide Manhattan plots for SNPs (grey), HLA alleles (red/green) and C4 haplotypes (blue) in (a) PGC MDD studies, (b) UK Biobank, and (c) meta-analysis of PGC MDD studies and UK Biobank.

In the UKB – PGC-MDD meta-analysis, no HLA allele or C4 haplotype met region-wide significance, however, HLA-B*0801 met the candidate threshold (p = 1.26e-4, OR = 0.98, 95% CI = 0.97-0.99) (Figure 1c). 143 SNPs reached region-wide significance (Supplementary Table 4). The variant with the lowest p-value (“metaanalysis index variant”) was a SNP in the classical class I region; rs9262120 (p = 8.74e-8, OR = 1.03, 95% CI = 1.02-1.05). The meta-analysis index SNP was in LD with the other 142 significant variants (0.44 < R^2^ < 1.00), and with the most significant SNP within the MHC in the PGC MDD GWAS^24^ (rs115507122, named rs3095337 in this study, R^2^ = 0.66), which did not reach region-wide significance in the meta-analysis (p = 7.14e-6, OR = 1.03, 95% CI = 1.01-1.04).

We identified six autoimmune diseases with evidence for a bi-directional relationship with depression: Crohn’s disease, multiple sclerosis, primary adrenocortical insufficiency, psoriasis vulgaris, systemic lupus erythematosus (SLE) and type 1 diabetes mellitus^9,13^. We identified 15 HLA alleles associated with risk for these autoimmune diseases (p < 3.9e-6) in European populations^46–57^, with a MAF > 0.05 in our study. Three HLA alleles had evidence for association with depression after correcting for multiple testing (p < 0.003): HLA-B*08:01 (for psoriasis and SLE), HLA-DQB1*02:01 (SLE) and HLA-DRB1*03:01 (multiple sclerosis, primary adrenocortical insufficiency, SLE) (Table 2).

**Table 2:** HLA alleles associated with risk for six autoimmune diseases and their association with depression in the PGC, UKB and meta-analysis. Bolded p-values met correction for multiple testing. Frq: allele frequency. OR: odds ratio.

| Trait | HLA allele | PGC |  | UKB |  | META-ANALYSIS |  |  |
| --- | --- | --- | --- | --- | --- | --- | --- | --- |
|  |  | Frq | OR | Frq | OR | OR | 95% CI | p-value |
| Crohn's disease<sup>46,47</sup> | HLA-A*03:01 | 0.15 | 0.96 | 0.14 | 0.99 | 0.99 | 0.98-1.00 | 0.176 |
|  | HLA-C*06:02 | 0.09 | 1.00 | 0.09 | 1.02 | 1.02 | 1.00-1.04 | 0.043 |
|  | HLA-DRB1*07:01 | 0.13 | 1.01 | 0.14 | 1.01 | 1.01 | 1.00-1.02 | 0.179 |
|  | HLA-DRB1*13:02 | 0.05 | 0.97 | 0.04 | 0.99 | 0.99 | 0.97-1.01 | 0.431 |
| Multiple sclerosis<sup>48,49</sup> | HLA-DQB1*03:02 | 0.11 | 1.00 | 0.10 | 1.00 | 1.00 | 0.99-1.01 | 0.537 |
|  | HLA-DRB1*03:01 | 0.13 | 0.95 | 0.15 | 0.98 | 0.98 | 0.97-0.99 | **0.003** |
|  | HLA-DRB1*15:01 | 0.14 | 0.98 | 0.14 | 1.00 | 0.99 | 0.98-1.00 | 0.355 |
| Primary adrenocortical insufficiency (Addison's disease)<sup>50,51</sup> | HLA-DRB1*03:01 | 0.13 | 0.95 | 0.15 | 0.98 | 0.98 | 0.97-0.99 | **0.003** |
| Psoriasis vulgaris<sup>52,53</sup> | HLA-A*02:01 | 0.28 | 1.01 | 0.27 | 1.01 | 1.01 | 1.00-1.02 | 0.005 |
|  | HLA-B*08:01 | 0.12 | 0.96 | 0.14 | 0.98 | 0.98 | 0.97-0.99 | **1.26e-4** |
|  | HLA-C*06:02 | 0.09 | 1.00 | 0.09 | 1.02 | 1.02 | 1.00-1.04 | 0.043 |
|  | HLA-DQA1*02:01 | 0.13 | 1.01 | 0.14 | 1.01 | 1.01 | 1.00-1.02 | 0.212 |
| Systemic lupus erythematosus<sup>54,55</sup> | HLA-B*08:01 | 0.12 | 0.96 | 0.14 | 0.98 | 0.98 | 0.97-0.99 | **1.26e-4** |
|  | HLA-DQA1*01:02 | 0.20 | 0.97 | 0.19 | 0.99 | 0.99 | 0.98-1.00 | 0.163 |
|  | HLA-DQB1*02:01 | 0.13 | 0.95 | 0.15 | 0.98 | 0.98 | 0.97-0.99 | **0.002** |
|  | HLA-DRB1*03:01 | 0.13 | 0.95 | 0.15 | 0.98 | 0.98 | 0.97-0.99 | **0.003** |
| Type 1 diabetes | HLA-A*24:02 | 0.08 | 0.97 | 0.07 | 1.01 | 1.00 | 0.98-1.02 | 0.578 |
| mellitus<sup>56,57</sup> | HLA-DPB1*01:01 | 0.05 | 0.92 | 0.06 | 0.99 | 0.98 | 0.96-1.00 | 0.067 |
|  | HLA-DPB1*04:02 | 0.12 | 0.99 | 0.11 | 1.00 | 1.00 | 0.99-1.01 | 0.857 |

Genetically predicted C4A brain expression was not significantly associated with depression status in the PGC (p = 0.066, OR = 1.06, 95% CI = 1.00-1.13), the UKB (p = 0.333, OR = 1.01, 95% CI = 0.99-1.03), or the meta-analysis (p = 0.150, OR = 1.01, 95% CI = 0.99-1.03). Evidence for association with HLA-B*08:01 remained after conditioning on genetically predicted C4A brain expression with the p-value decreasing from p = 1.26e-4 to p = 3.39e-5 in the conditional analyses. The meta-analysis results for genetically predicted C4A brain expression and four common C4 haplotypes are summarised in Figure 2. The full results for HLA alleles, C4 haplotypes and conditional analyses in the PGC, UKB and metaanalysis are provided in Supplementary Table 5.

**Figure 2:**
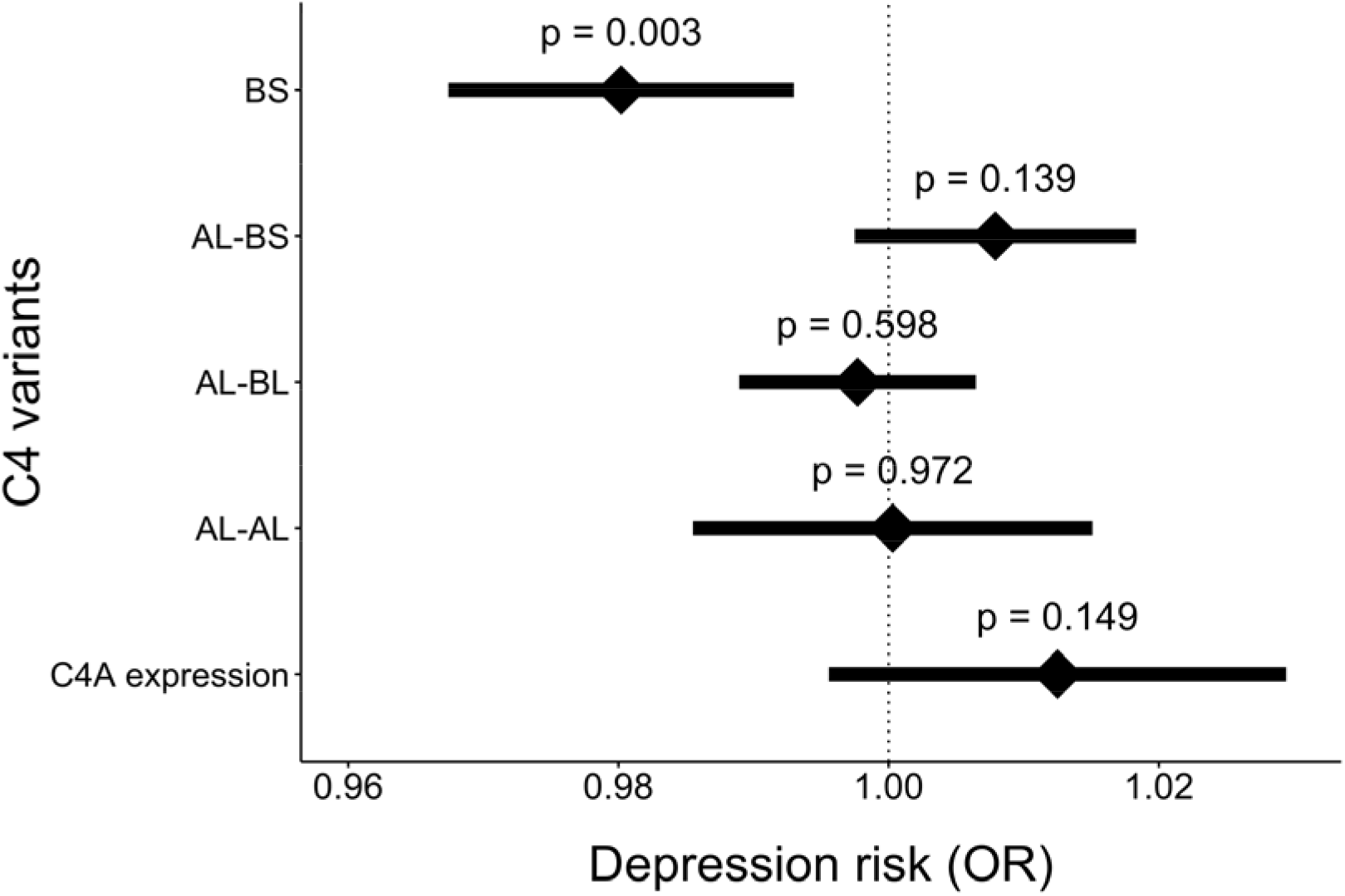
Association of genetically predicted C4A brain expression and four C4 haplotypes in the metaanalysis of PGC-MDD studies and UK Biobank. Error bars show 95% confidence intervals.

## Discussion

To further understand the mechanisms driving comorbid autoimmunity and depression, we investigated evidence for shared genetic influences in the MHC, a region harboring genetic risk for autoimmune diseases and psychiatric disorders. Our primary aim was to test HLA alleles and C4 haplotypes for association with depression. Under a conservative region-wide significance threshold, we found no evidence that HLA alleles, which play a major role in susceptibility to autoimmune diseases, or C4 haplotypes, which associate strongly with risk for schizophrenia, also confer risk for depression. However, under a candidate threshold, HLA-B*0801 had significant evidence for association with depression status. Conditioning on genetically predicted C4A brain expression did not diminish signal from HLA-B*0801, indicating independence from C4 haplotypes.

We further explored common HLA alleles associated with autoimmune diseases that have evidence of a bi-directional relationship with depression. The strongest evidence for association with depression was HLA-B*08:01, followed by HLA-DQB1*02:01 and HLA-DRB1*03:01. Previous studies have shown that all three HLA alleles are risk increasing for SLE^54,55^, HLA-DRB1 *03:01 is also risk increasing for multiple sclerosis^48,49^ and primary adrenocortical insufficiency^50,51^, and HLA-B*08:01 is also risk increasing for psoriasis^52,53^. In contrast, our findings indicate that HLA-B*08:01, HLA-DQB1*02:01 and HLA-DRB1*03:01 have modest protective effects in depression, indicating these alleles do not harbor shared risk for autoimmune disease and depression.

Imputation of C4 haplotypes identified four common haplotypes, none of which were associated with risk for depression in the PGC studies, in the UKB or the meta-analysis. These results are in stark contrast to schizophrenia where association with C4 haplotypes accounts for most of the observed SNP association in the HLA region. Our results suggest the C4 association does not contribute to the common genetic susceptibility between depression and schizophrenia, observed in a genetic correlation of rG=0.34 (p = 7.7e-40) between these disorders^24^.

At the level of region-wide significance, we detected 70 SNPs associated with depression in the UKB sample, and 143 in the meta-analysis. In each case, the top SNP was in moderate to strong LD with other significant variants, indicating a single peak of independent association. We found consistency in SNP signal between our study and the PGC-MDD GWAS of depression^24^, with the top SNPs in each study showing moderate to strong LD. This was not unexpected given that our study is a subset of the studies included in the PGC-MDD meta-analysis^24^.

The true identity of causal variants within the MHC remains unresolved, and fine-mapping within the MHC is challenging due to the high density of genetic variation and strong LD. Our results strongly suggest that the association signal observed in the MHC in depression^24,25^ does not arise from HLA alleles or C4 haplotypes. These results suggest that any associations with these variants are either rare or have very modest effect sizes. We note that Howard, et al.^25^ increased power by leveraging a broader phenotyping approach. It is interesting to speculate that the broader depression phenotype captures individuals distressed by physical disease. This would go some way to explaining signal in the MHC, which has evidence for association with more diseases than another other region of the genome^20^. However, a more parsimonious explanation could be that MHC signal in depression maps to SNPs or to other genetic loci not imputed in this study. This is highly plausible in light of the fact that the MHC contains more genes than any other region in the human genome^20^. Under this scenario, large sample sizes and sequencing may be required to dissect SNP signal within the MHC.

Although our findings do not support a role for HLA alleles within the MHC in risk for depression, it is possible that shared genetic risk for depression and autoimmune diseases is situated outside the MHC. Efforts to identify genome-wide pleiotropy were undertaken in the recent PGC-MDD GWAS using LD Score to estimate genetic correlations between depression and several autoimmune diseases^24^. There was no evidence for significant cross-trait correlations; the strongest observed was between depression and inflammatory bowel disease (rG = 0.07, p = 0.06). In other efforts to detect genome-wide pleiotropy, Euesden et al.^14^ found no evidence that polygenic risk scores (PRS) for rheumatoid arthritis predicted depression status in an independent sample, nor did PRS for depression predict autoimmune disease status.

One possibility is that there is a sub-group of individuals enriched for depression and autoimmune risk alleles. Under this scenario, there may be insufficient power to detect the relationship. Identifying, for example, a sub-group of individuals with depression, who are also enriched for autoimmune risk alleles, would go some way to explaining the observed comorbidity between these traits. Furthermore, identifying a sub-type of depression, e.g. an ‘immune related’ depression group, would help to dissect heterogeneity in the depression phenotype.

In summary, this study is the first to interrogate the involvement of HLA alleles and C4 haplotypes in depression risk, and we find no evidence that either type of genetic variant plays a major role in susceptibility for depression. In contrast, the three HLA alleles that showed nominal significance in our study conferred modest protective effects for depression. Furthermore, the strong association for C4 alleles seen in schizophrenia is absent in depression. Large sample sizes and regional sequence data may be required to dissect SNP signal within the MHC.

## Acknowledgements

This work was supported by a PhD studentship awarded to K.P.G. from the UK Medical Research Council. PFO receives funding from the UK Medical Research Council (MR/N015746/1).

This paper represents independent research part-funded by the National Institute for Health Research (NIHR) Biomedical Research Centre at South London and Maudsley NHS Foundation Trust and King’s College London. The views expressed are those of the author(s) and not necessarily those of the NHS, the NIHR or the Department of Health and Social Care.

We thank participants and scientists involved in making the UK Biobank resource available (http://www.ukbiobank.ac.uk/). UK Biobank data used in this study were obtained under approved application 18177.

We are deeply indebted to the investigators who comprise the PGC, and to the hundreds of thousands of subjects who have shared their life experiences with PGC investigators. The PGC has received major funding from the US National Institute of Mental Health and the US National Institute of Drug Abuse (U01 MH109528 and U01 MH1095320). Acknowledgements and funding sources for each of the primary studies in the PGC are contained in supplementary materials.

Statistical analyses were carried out on the NL Genetic Cluster Computer (http://www.geneticcluster.org/) hosted by SURFsara, and the King’s Health Partners High Performance Compute Cluster funded with capital equipment grants from the GSTT Charity (TR130505) and Maudsley Charity (980).

## Previous presentation of this data

K.P.G presented a poster entitled “Genetic variation in the Major Histocompatibility Complex and association with depression” at the World Congress of Psychiatric Genetics in Glasgow, UK on 14th October 2018.

